# Emodin Attenuates Renal Interstitial Fibrosis via Regulation of TIMP1/MM9 Pathway in Rats

**DOI:** 10.1101/736876

**Authors:** Dongmei Li, Qingchuan Zhang, Yujiao Lou, Chunfeng He, Renze Gu, Wei Liu

**Affiliations:** Putuo Hospital, Shanghai University of Traditional Chinese Medicine, Shanghai 200333, China; The Ninth People’s Hospital of Shanghai, Shanghai Jiao Tong University, Shanghai, 200030, China

**Keywords:** Emodin, MMP9, TIMP1, UUO, Renal Interstitial Fibrosis

## Abstract

Emodin has a variety of pharmacological functions including anti-bacterial infection, anti-inflammatory, anti-oxidation, anti-tumor, regulation of gastrointestinal activities and anti-hepatic and lung fibrosis. However, the role of emodin in the regulation of renal interstitial fibrosis(RIF) remains poorly understood. In this study, we investigated the regulation of emodin in RIF and revealed the underlying molecular mechanisms. We established a unilateral ureter obstruction(UUO) model to simulate renal interstitial fibrosis in rats. We found that UUO rats were observed a large amount of inflammatory cell infiltration, more fibroblast proliferation, collagen fiber proliferation. Furthermore, we demonstrated that the up-regulation of TIMP1 and down-regulation of MMP9 were related to renal fibrosis. However, those phenomena in emodin-treated UUO rats were ameliorated. Collectively, our results provide new insights into the treatment of RIF and suggest that the TIMP1/MM9 signaling axis may be a potential therapeutic target for RIF.

## Introduction

Renal Interstitial Fibrosis(RIF) is a hallmark of pathological progression from to renal injuries to a variety of Chronic Kidney Disease(CDK)[1, 2]. CKD affects over 10% of the worldwide population with high mortality, due to limited available or affordable treatment partly. What’s more, the number will further increase according to the increased number of elderly people and increased prevalence of major disease which leads to CKD like obesity, diabetes, atherosclerosis and hypertension[3–5]. Because RIF is considered as the underlying pathological process of chronic CKD, therefore, it is necessary to discover and identify the novel and specific treatment to inhibit fibrosis and ameliorates CDK.

Fibrosis is defined by excessive accumulation of extracellular matrix(ECM) mainly produced by tissue resident mesenchymal cells, accompany with tissue regeneration and inflammation. In normal kidney, the production and degradation of ECM are dynamic and in balance, but the balance is broken in fibrotic ECM[2]. The ECM is controlled by matrix metalloproteinases (MMPs) and tissue inhibitors of metalloproteinases (TIMPs). MMPs are the major enzymes implicated in ECM degradation. It is reported that four forms of TIMPs all are expressed in kidney, but only TIMP1/2 inhibits MMPs expression. MMP9 is an important member of the MMPs family. It participates in the degradation of ECM, regulation the synthesis and metabolic balance of ECM under physiological and pathological conditions. TIMP1 specifically blocks MMP-9 activity, which plays a key role in ECM accumulation and degradation while TIMP-2 involves in regulation of MMP-2 activity[6–8].

Emodin is an anthraquinone compound extracted from rhubarb and has a wide range of pharmacological effects. In liver fibrosis, recent studies showed that Emodin alleviates CCl4-induced by suppressing EMT and transforming growth factor-ß1 in rats and it reduced infiltration of Gr1hi Monocytes[9, 10]. In bleomycin-Induced pulmonary fibrosis, it was also reported that emodin attenuates fibrosis by suppressing EMT, fibroblast activation, anti-inflammatory and anti-oxidative activities in Rats[11, 12]. Although emodin plays certain positive therapeutic effects on liver and lung fibrosis, the underlying pharmacology and mechanism on renal fibrosis remain unclear. Because emodin cloud inhibit expression of both TGF-ß1 and TIMP-1 in an immortalized rat hepatic stellate cell line[13].

Here, we established a unilateral ureteral obstruction model of rats and performing HE or MASSON staining. IHC observation of the expression of positive and negative regulatory molecules TIMP1 and MMP9 were also performed to provide the mechanism of renal fibrosis. We wonder whether emodin also plays the anti-fibrosis role in renal fibrosis, aiming to provide a theoretical basis for clinical prevention and development of renal fibrosis.

## Materials and Methods

### UUO Experimental Animal Model

All animal experiments were performed according to the Putuo Hospital, Shanghai University of Traditional Chinese Medicine for the Care and Use of Laboratory Animals. Male Wister rats (180-200g) were obtained from Shanghai Slrk Laboratory Animal(Shanghai, China). Experimental Unilateral Ureter Obstruction (UUO) represents a model for obstructive nephropathy but also allows insight into the process of interstitial fibrosis that is a common characteristic of many chronic nephropathies. In this model, the ureter was separated from surrounding tissues and the left ureter was ligatured. Animals were randomly divided into the following groups: non-surgery, UUO alone or UUO with Emodin treatment. UUO alone and UUO with Emodin groups were subject to UUO on Day 0 and the kidneys are removed on day 3, 7, 14 and 21.

### RNA-seq and gene expression detection

Total RNA was isolated from each thymic sample using the standard TRIzol protocol (Invitrogen, Carlsbad, CA, USA). RNA quality was examined by gel electrophoresis and with a Nanodrop spectrophotometer (Thermo, Waltham, MA, USA). High-throughput mRNA sequencing (RNA-seq) was carried out using Illumina HiSeq X Ten instrument. Qualified reads were mapped to the human hg19 reference genome (hg19) at the same time by TopHat (v2.1.0) with default parameters. In this study, bioinformation analysis was mainly performed by using R language(https://www.r-project.org/) with several publicly available packages. A probability value (p) < 0.05 was considered to be significant in this study.

### Western Blot

Total proteins were extracted from the kidneys was measured by BCA protein assay kit (Beyotime, Shanghai, China). Western blot assay was performed with the standard method. Anti-MMP9 (1:500, Abcam, Cambridge, UK, Cat: 76003) and anti-TIMP1 (1:500, Abcam, Cambridge, UK, Cat: ab61224) were used as the primary antibodies.

### Immunohistochemistry

Tissues were fixed with 4% paraformaldehyde for at least 24 hours and then they were dehydrated, embedded and cut into 4μm sections. The sections were stained with haematoxylin & eosin(H&E) and MASSON staining for general morphological assay. Immunohistochemical detection was conducted with general approaches. Anti-MMP9 (1:500, Abcam, Cambridge, UK, Cat: 76003) and anti-TIMP1 (1:500, Abcam, Cambridge, UK, Cat: ab61224) were used as the primary antibodies.

### Statistical Analyses

The data are presented as mean ± SD in the figures. A Student t test was performed to compare the in vitro data. P values less than 0.05 were considered statistically significant.

## Results

### Emodin ameliorates renal interstitial damage in UUO Rats

To explore the function of emodin on renal interstitial fibrosis, we established UUO model rats. H&E staining were performed to observe the structure in UUO and emodin-treated UUO groups compare with normal renal. In the UUO group, renal tubular epithelial cell atrophy, brush border disappeared, lumen collapse and cystic lumen were observed comparing with the normal group. Those features were ameliorated in emodin-treated UUO group (Figure 1A). Consistent with the changes in the H&E staining, Masson’s trichrome staining showed that the Emodin-treated UUO group exhibited a marked reduction in collagen deposition (Figure 1B). These results demonstrate that emodin ameliorates renal interstitial damage in UUO Rats.

**Figure 1.**
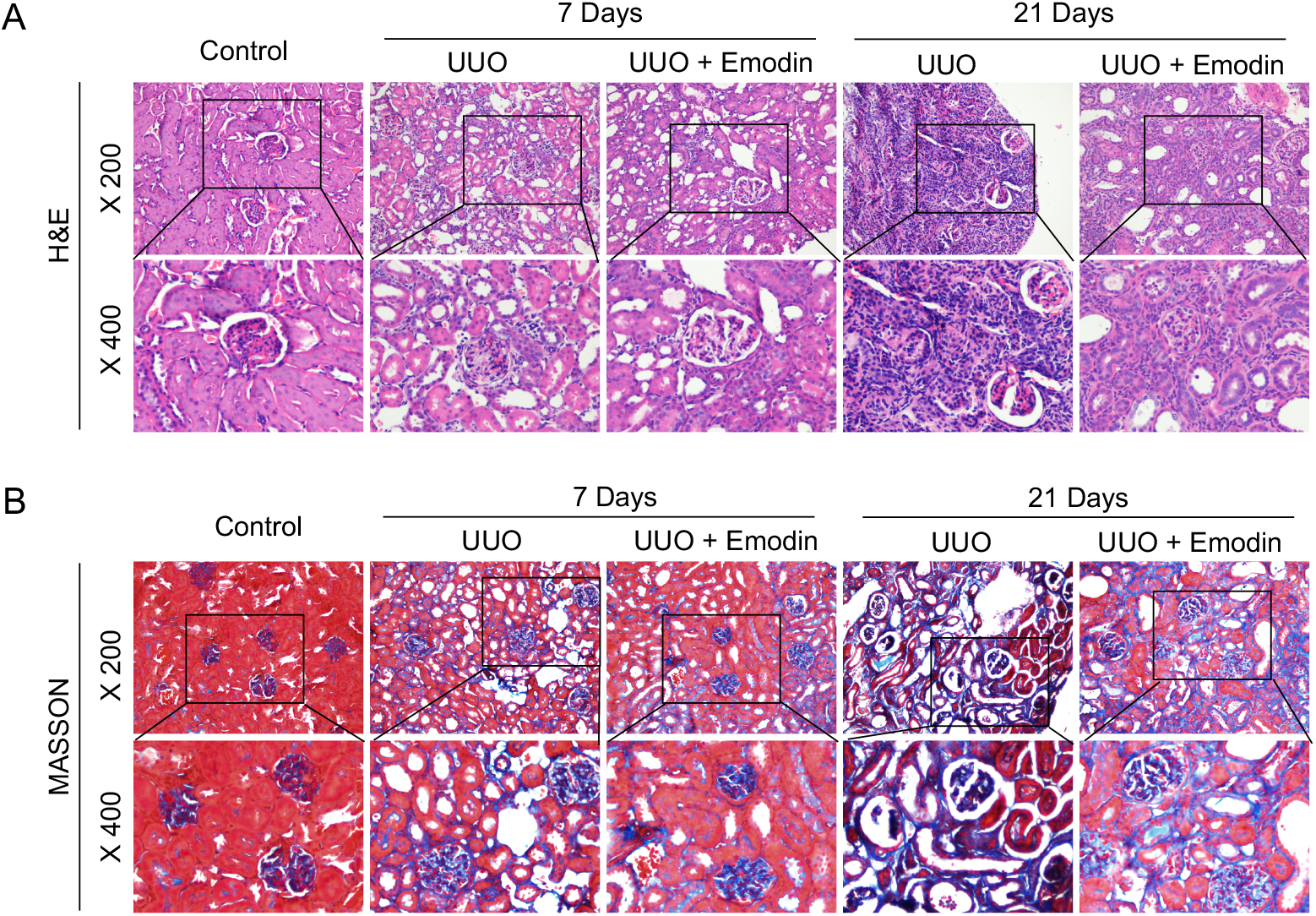
Effect of emodin in renal fibrosis induced by unilateral ureteral obstruction (UUO). A, Representative H&E staining images of kidney tissue 7 days and 21 days after UUO. B, Representative MASSON staining images of kidney tissue 7 days and 21 days after UUO.

### Emodin inhibits the expression of TIMP1 while up-regulates MMP9

Emodin demonstrated the function of attenuateing renal interstitial fibrosis from the analysis above. To explore the mechanism of Emodin, we identified 175 (66 up-regulated and 109 down-regulated) differentially expressed genes between UUO treated with emodin group and UUO alone group(Figure 2A and 2B). The GO enrichment analysis showed that based on these genes were significantly enriched in the excretion, dendritic spine membrane, water channel activity(Figure2C). Besides, the hallmark gene sets analysis found fatty acid metabolism and TNFα signaling via NF-kB were enriched in emodin treated group(Figure 2D) while allograft rejection and inflammatory response were negatively correlated with emodin(Figure 2E). In addition, The RNA-seq showed that the group treated by emodin influenced the expression of MMPs family and TGF family (Figure 2F and 2G). It has been reported that the myofibroblasts are contractile and augment the deposition of the extracellular matrix, which leads to progressive interstitial fibrosis. This process is enhanced by the decrease of ECM degradation[14]. MMP9 is of one the important enzyme which could degrade extracellular matrix, which is regulated by TIMP1[15–17], so we detected the expression of MMP9 and TIPM1 in UUO model. It was shown that the low MMP9 expression but high TIMP1 expression in UUO group while an increased MMP9 expression but decreased TIMP1 expression in emodin-treated UUO group (Figure 3A). Together, the result suggests that emodin inhibits the expression of TIMP1 while up-regulates MMP9.

**Figure 2.**
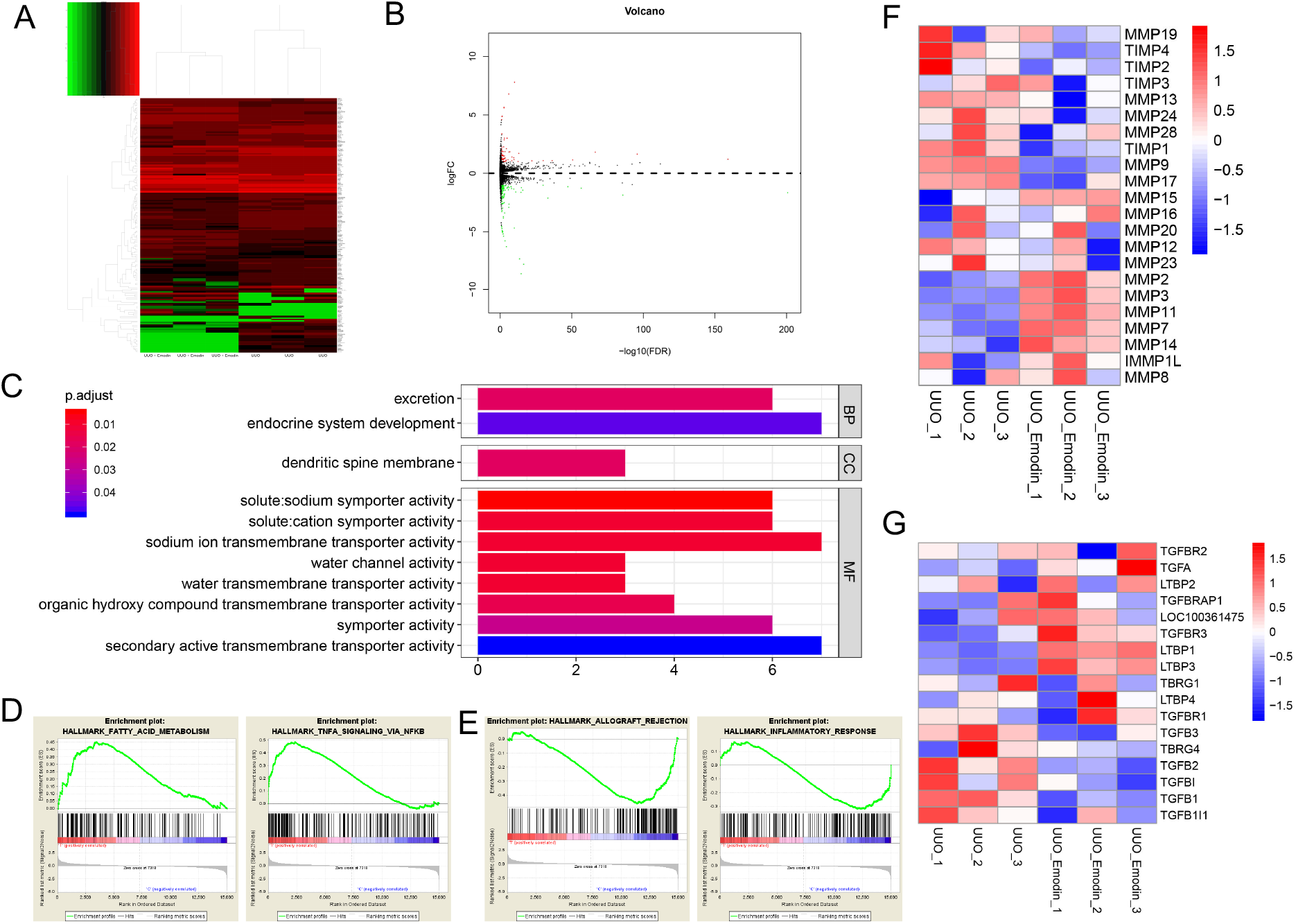
Effect of emodin on the regulation of TIMP1 and MMP9 in UUO rats. Representative images of TIMP1 and MMP9 Immunohistochemistry staining of kidney tissue 7 days after UUO.

**Figure 3.**
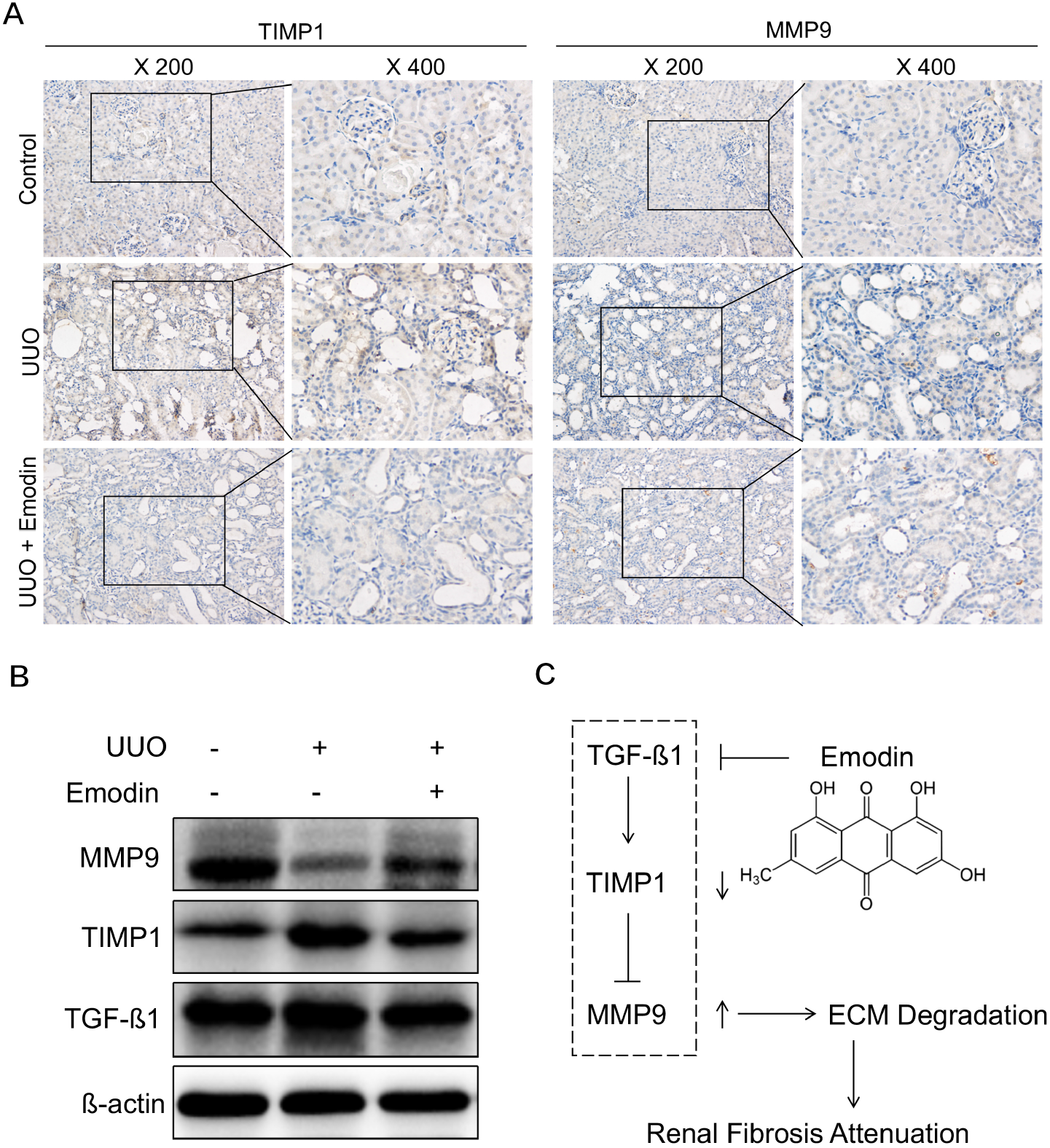
Effect of emodin on the regulation of TGF-β1 in UUO rats. A, Immunoblotting of TGF-β1, TIMP1 and MMP9 in UUO and emodin-treated UUO groups comparing with normal group. B, The schematic model of the role of emodin in regulating renal fibrosis.

### Emodin ameliorates renal fibrosis in rats via TGF-ß1 pathway

Recent evidences have shown that TGF-ß1 are associated with ECM accumulation in organ fibrosis, suggesting the existence of a link between TGF-ß1 and fibrosis[18]. We hereby tested the effect of the emodin on TGF-ß1 and found that emodin-treated group down-regulated the expression of TGF-ß1. Moreover, consistent with the Immunohistochemistry staining results ahead, it showed that emodin ameliorated renal fibrosis by inhibiting the expression of TGF-ß1 and TIMP1 while up-regulates MMP9 (Figure 3B and 3C).

## Discussion

Recent studies have demonstrated that emodin is associated with anti-inflammatory, anti-infection, anti-oxidation and regulation of gastrointestinal activities. Previous studies showed that emodin are effective in the treatment of pulmonary fibrosis through notch signaling pathway. However, the functions of emodin and its underlying molecular mechanisms as a fibrosis suppressor in renal fibrosis remain unclear. Our experiment verified emodin influence on anti-fibrosis in renal and found another pathway involved. In our study, we have revealed the function of emodin in weakening TIMP1-regulated MMP9 inhibition and illustrated the underlying molecular mechanisms involved with TGF-β1.

Renal Interstitial Fibrosis is regarded as a common pathological progression to a variety of Chronic Kidney Disease. According to model of unilateral ureteral obstruction (UUO) in the rodent could generate progressive renal fibrosis[19], we used UUO model to study the effect of emodin on renal fibrosis. Because fibrosis is closely related with ECM accumulation and decreased despot ion and this process is control by the enzymatic actions of the decreased secreting of proteases matrix metalloprotease(MMP), which are regulated by tissue inhibitors of metalloproteases (TIMP). Recent report also shown that emodin effectively inhibited PMA and TGF-β1stimulated TIMP-1 expression in hepatic stellate cells. TGF-β1 has been proved to be the up-stream of TIMP1. It inhibits the degradation of extracellular matrix by decreasing the synthesis of MMPs and increasing the synthesis of inhibitors of MMPs. So we first used MMP9 and TIMP1 as our detection marker to estimate the effect of emodin in renal fibrosis and then detected their up-stream regulator TGF-β1.

Based on the above observation, we used UUO model rats. We found that the emodin-treated UUO group ameliorated renal interstitial damage compared with UUO Rats. Emodin-treated UUO group showed a significant reduction in collagen deposition and fibroblast proliferation. The expression of MMP9 was increased in emodin-treated UUO while the expression of TIMP1 was decreased. Together, the result suggests that emodin inhibits the expression of TIMP1 while up-regulates MMP9. TGF-ß1 was also inhibited in emodin-treated group. That is to say, emodin may ameliorate renal fibrosis by inhibiting the expression of TGF-ß1 and TIMP1 while up-regulates MMP9.

In summary, emodin showed an anti-fibrosis effect in UUO renal fibrosis model. It suggests that emodin may act as a promising candidate to slowdown renal fibrosis. However, the clinical use of emodin on renal fibrosis should be further investigated.

## Acknowledgement

This work was supported by the funds from Putuo Hospital, Shanghai University of Traditional Chinese Medicine Foundation (China, Grant/Award Number: 2016324A). And also supported by the funds from Shanghai University of Traditional Chinese Medicine Foundation(Grant/Award Number: 18Lk060).

